# Analytical strength-duration curve for the spiking response of the LIF neuron to an alpha-function-shaped excitatory current pulse

**DOI:** 10.1101/2023.09.05.556358

**Authors:** Alexander Paraskevov

## Abstract

Whether or not the neuron emits a spike in response to stimulation by an excitatory current pulse is determined by a strength-duration curve (SDC) for the pulse parameters. The SDC is a dependence of the minimal pulse amplitude required to elicit the spiking response on either the pulse duration or its decay time. Excitatory neurons affect the others through pulses of excitatory postsynaptic current. A simple yet plausible approximation for the time course of such a pulse is the alpha function, with linear rise at the start and exponential decay at the end. However, an exact analytical SDC for this case is hitherto not known, even for the leaky integrate-and-fire (LIF) neuron, the simplest spiking neuron model used in practice. We have obtained general SDC equations for the LIF neuron. Using the Lambert W function — a widely-implemented special function, we have found the exact analytical SDC for the spiking response of the LIF neuron stimulated by an excitatory current pulse in the form of the alpha function. To compare results in a unified way, we have also derived the analytical SDCs for (i) rectangular pulse, (ii) ascending ramp pulse, and (iii) instantly rising and exponentially decaying pulse. In the limit of no leakage, we show that the SDC is reduced to the classical hyperbola for all considered cases.

## 1. Introduction

For modeling biological neuronal networks, an adequate choice of the amplitude and duration values for pulses of excitatory postsynaptic current (EPSC) is evidently crucial [1–3]. No less important is the waveform – or shape – of such a pulse. The same can be said for the stimulating electrical pulses in precision neuroprosthetics [4–7]. Whether or not the neuron emits a spike in response to an excitatory current pulse (synaptic or externally-stimulated via sealed microelectrode in the current-clamp mode of whole-cell patch-clamp technique) is determined by the so-called strength-duration curve (SDC) [8–14]. The SDC is a dependence of the minimal pulse amplitude required to elicit the spiking response on either the pulse duration or its decay time, provided that the neuron’s initial state is unaltered.

Leaky Integrate-and-Fire (LIF) neuron is the simplest dynamic model of a spiking neuron that is widely used in theoretical studies and large-scale simulations of spiking activity in biological neuronal networks [13]. The analytical SDC for the LIF neuron stimulated by a rectangular current pulse was derived by Lapicque just after the seminal LIF model definition [15] (see also other early theoretical findings [16–19]).

Despite its analytical tractability, the rectangular pulse is poorly suitable both for modeling an EPSC pulse and for artificial pulse stimulation of biological neurons in neuroprosthetics [4]. In turn, the alpha function, with the linear rise at the start and exponential decay at the end (see the inset on the right graph in Fig. 1), is a simple yet plausible approximation for the time course of an EPSC pulse caused by the activation of AMPA receptors [11, 20, 21]. However, an analytical SDC for the spiking response of the LIF neuron stimulated by an excitatory current pulse in the form of the alpha function has been hitherto unknown.

**FIG. 1.**
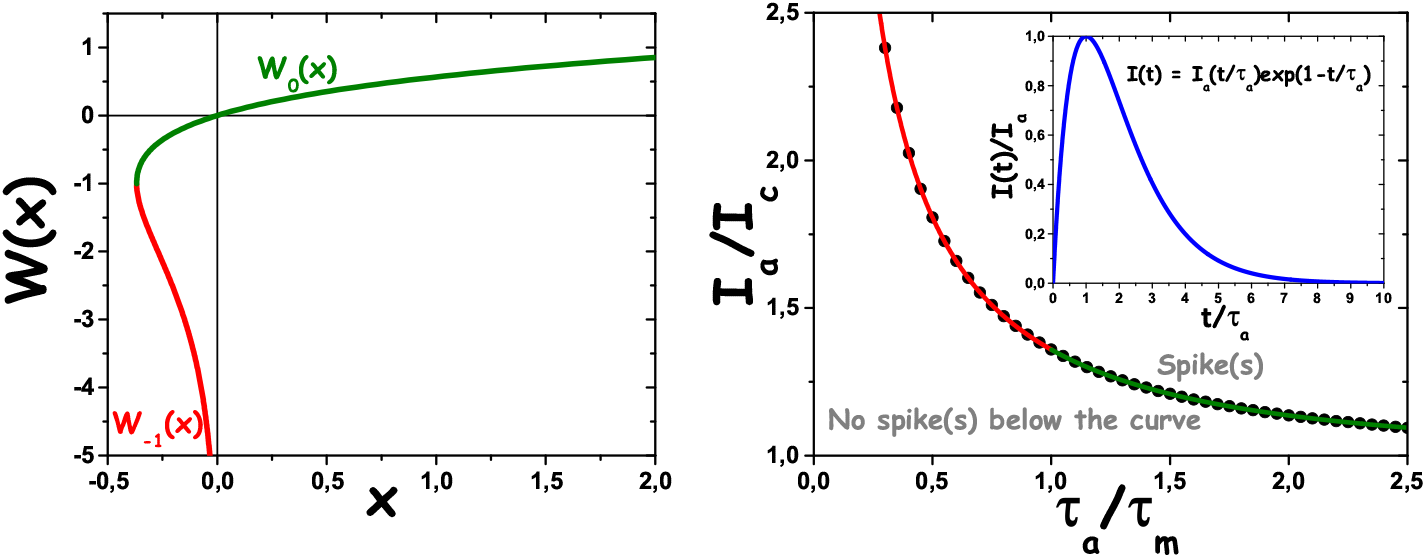
Left graph: The Lambert function *W* (*x*) with its two branches highlighted and standardly denoted. Right graph: The solid line composed of the red and green parts is an analytically derived strength-duration curve (SDC) for the LIF neuron stimulated by a single excitatory current pulse in the form of the alpha function shown in the inset. The red and green parts of the SDC correspond to the underlying branches of *W* (*x*) (see Eqs. (24) and (25)). The numerically calculated SDC is shown by filled black circles. It accurately matches the analytical SDC. Parameters *I*_*c*_ = 150 pA and *τ*_*m*_ = 20 ms are constants of the LIF neuron model (see Sec. 2).

In this paper, we have shown that the exact analytical SDC (Fig. 1, right graph) for the above case can be expressed through the branches of the special Lambert W function [22, 23] (Fig. 1, left graph) that gives the solution *y* = *W* (*x*) of transcendental equation *y* exp(*y*) = *x*. The SDC depends on two compound parameters of the LIF model: minimal ‘critical’ (or ‘rheobase’) direct current *I*_*c*_ for periodic spike generation and the ‘membrane’ relaxation time constant *τ*_*m*_. Notably, these are standardly accessible in patch-clamp experiments.

We assume that the obtained result can be used for tuning the parameters of network models with LIF neurons [24–28], and for calibrating the current-based stimulation of a single neuron in electrophysiological experiments. In addition, these results can be used in theoretical studies of synaptic plasticity, e.g., for analyzing the influence of spike-timing-dependent plasticity (STDP) on integrative properties of the LIF neuron. In particular, the multiplicative version of the classical pair-based STDP rule [29] explicitly contains a maximal synaptic weight; it is also implicitly introduced in the additive version. A difficult question for a modeler of biological neuronal networks is that from what considerations one should take the maximal weight value, implying that the synaptic weight is either the EPSC amplitude or EPSC upward slope. One can suggest determining the maximal weight value from the analytical SDC for the EPSC pulse, provided that its typical duration is known. Indeed, any higher weight value is hardly meaningful while considering integrative excitation of the postsynaptic neuron: having such a ‘strong’ incoming synapse activated, the neuron will generate a spike regardless of other incoming excitatory signals, provided that inhibitory signals do not come for a while.

## 2. Standard LIF neuron model

Subthreshold dynamics of transmembrane potential *V* for the LIF neuron is described by equation

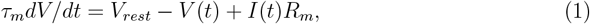

where *V*_*rest*_ is the neuron’s resting potential, *τ*_*m*_ = *C*_*m*_*R*_*m*_ is the characteristic time for relaxation of *V* (*t*) to *V*_*rest*_, *C*_*m*_ and *R*_*m*_ are the capacity and resistance of the neuron’s membrane, *I*(*t*) is the total incoming synaptic current, which, as a function of time *t*, depends on the choice of the synapse dynamic model and the number of incoming synapses. When the transmembrane potential reaches a threshold value *V*_*th*_ = *V* (*t*_*sp*_), it is supposed that the neuron emits a spike, then *V* abruptly drops to *V*_*rest*_ and retains this value during the absolute refractory period *τ*_*ref*_,

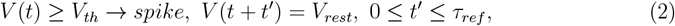

and afterwards the potential dynamics is again described by Eq. (1). The result of the LIF neuron dynamics is a sequence of spike generation moments 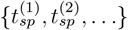.

If the total incoming current in Eq. (1) is a positive constant, *I*(*t*) = *I* > 0, and exceeds ‘rheobase’ or ‘critical’ value *I*_*c*_ = (*V*_*th*_−*V*_*rest*_)*/R*_*m*_, then the LIF neuron emits spikes regularly, with period

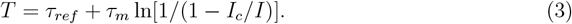

Finally, the numerical values of parameters for the LIF neuron are as follows: *τ*_*m*_ = 20 ms, *C*_*m*_ = 200 pF, *R*_*m*_ = 0.1 GΩ, *V*_*rest*_ = −65 mV, *V*_*th*_ = −50 mV. These give the critical current value *I*_*c*_ = 150 pA. A typical value for the refractory period *τ*_*ref*_ is 2 or 3 ms, though it will not be used further, as we study a single spiking response.

In addition, to make an accurate comparison with the earlier results, where the strength-duration curve was fitted by a hyperbola, it is also worth outlining the Perfect Integrate-and-Fire (PIF) neuron, which has no leakage current due to formally infinite resistance *R*_*m*_.

The PIF neuron dynamics is described by equation

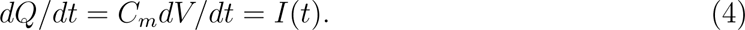

Here *Q* = *C*_*m*_(*V* − *V*_*rest*_) is excessive transmembrane electric charge, and all other notations are the same as those for the LIF neuron. Instead of the critical current *I*_*c*_, which is formally zero for the PIF neuron, it has the critical charge value *Q*_*c*_ = *C*_*m*_(*V*_*th*_ − *V*_*rest*_) at which the PIF neuron emits a spike. The after-spike dynamics for the PIF model is the same as for the LIF one, see Eq. (2). We used the same numerical values of corresponding parameters as for the LIF neuron, resulting in the critical charge value *Q*_*c*_ = 3 pC.

At last, suppose we have a formula derived for the LIF neuron, e.g., the period *T* of regular spiking, Eq. (3). How to transform the formula for describing the PIF neuron case? To this end, one should concurrently turn *τ*_*m*_ to infinity and *I*_*c*_ to zero such that their product *I*_*c*_*τ*_*m*_ = *Q*_*c*_ would hold a finite non-zero value, which is the critical charge. Following this way, the spiking period *T* for the PIF neuron stimulated by constant depolarizing current *I >* 0 is given by *T* = *τ*_*ref*_ + *Q*_*c*_*/I*.

## 3. General equations for the strength-duration curve (SDC)

By its definition, the SDC is a dependence of minimal spike-triggering amplitude *I*_*a*_ of the stimulation current pulse on its characteristic duration *τ*_*a*_. Assuming that the neuron is initially at rest, the corresponding evoked pulse of the membrane potential must have the amplitude equal to *V*_*th*_. In other words, the pulse maximum is reached at *V* (*t*_*sp*_) = *V*_*th*_, giving the second condition for SDC: *dV/dt* = 0 at *t* = *t*_*sp*_. Together, one gets the system of two algebraic equations

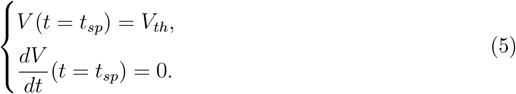

The solutions of the system are *I*_*a*_ and the moment *t*_*sp*_ of spike generation, as functions of *τ*_*a*_. The SDC is function *I*_*a*_(*τ*_*a*_).

For the LIF neuron, a straightforward integration of Eq. (1) with arbitrary current *I*(*t*) and the initial condition *V* (*t* = *t*_0_) = *V*_0_ results in

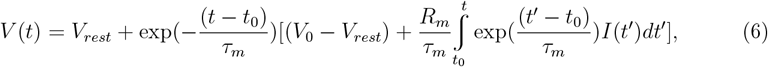

where *t*_0_ ≤ *t* ≤ *t*_*sp*_ and *t*_*sp*_ is the moment of the first spike since *t*_0_.

In what follows, we use *t*_0_ = 0 and *V*_0_ = *V*_*rest*_. Assuming *I*(*t*) being a stimulating pulse of amplitude *I*_*a*_ and characteristic decay time *τ*_*a*_, the system (5) is then as follows

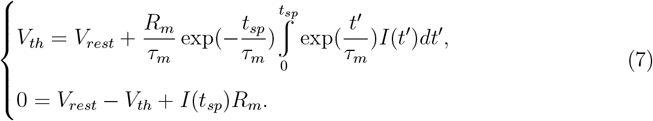

Using the standard notation for the critical current *I*_*c*_ = (*V*_*th*_ − *V*_*rest*_)*/R*_*m*_ and denoting *x* = *t*_*sp*_*/τ*_*m*_, we get

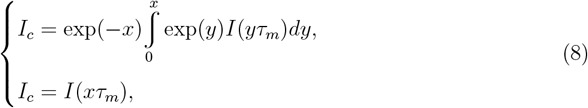

where *x* is the dimensionless moment of the first spike after the stimulation begins.

The Eqs. (8) do not work if the decay time of the stimulating pulse is zero, i.e., if the trailing edge of the pulse is artificially chopped off, like for the rectangular pulse or the ramp one (see Sec. 5). In this case, the first spike time *t*_*sp*_ is simply equal to the pulse duration.

## 4. SDC for the alpha-function-shaped pulse

Let us consider that the stimulation pulse *I*(*t*) has a shape determined by the alpha function,

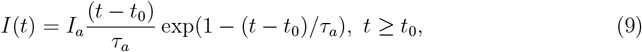

which is the solution of equations (where *δ*(*t* − *t*_0_) is the Dirac delta function) [30]

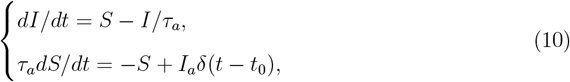

at *I*(*t* ≤ *t*_0_) = 0 and *S*(*t < t*_0_) = 0. Setting *t*_0_ = 0, we get Eqs. (8) in the form

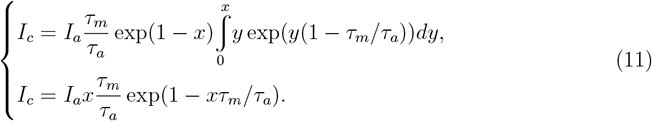

Denoting *k* = 1 − *τ*_*m*_*/τ*_*a*_ and, for *k* ≠ 0, integrating

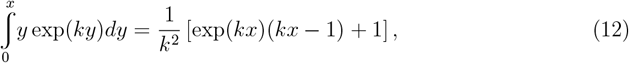

we get

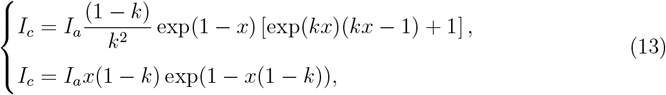

from where, combining both equations, it follows that

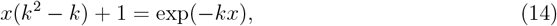

resulting in

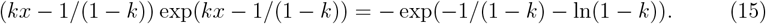

The latter equation can be directly solved using the Lambert W function definition: *y* = *W* (*b*) is the solution of *y* exp(*y*) = *b*, respecting the W function branches [22, 23]:

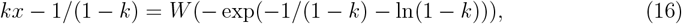

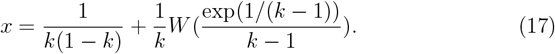

In turn, from the 2nd equation of system (13) one straightforwardly gets

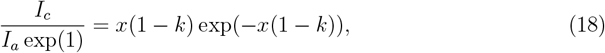

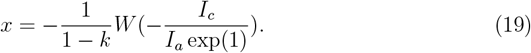

Comparing solutions (17) and (19), we get

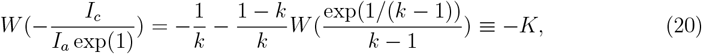

and, again using the definition of Lambert W function, only inside out now, we finally get

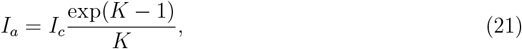

where

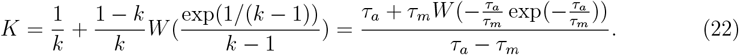

From Eq. (17) it follows that *K* = *t*_*sp*_*/τ*_*a*_, i.e., *K* is simply the 1st spike time, in units of *τ*_*a*_ (cp. [31, 32]).

If *k* = 0, i.e. *τ*_*a*_ = *τ*_*m*_, system (11) is reduced to

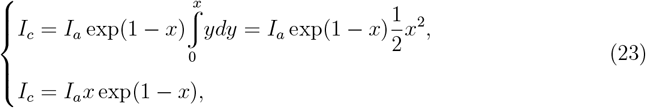

from where *x* = 2 and *I*_*a*_ = *I*_*c*_*/*(*x* exp(1 − *x*)) = *I*_*c*_ exp(1)*/*2. Notably, this result can be obtained from formula (21) by setting *K* = 2 there.

Taking into account the W function branches, the final formula for SDC is given by

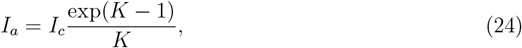

where *K* is the following function of *τ*_*a*_,

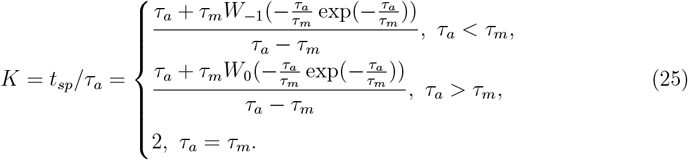

Here *W*_−1_(…) and *W*_0_(…) are the lower and upper branches of the Lambert W function (see Fig. 1, left graph), respectively.

Notably, at *τ*_*a*_ → +∞ one gets *K* → 1 in Eq. (25) and, according to Eq. (24),

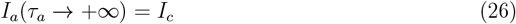

that is a general limiting case for the SDC, regardless of a specific shape of the stimulation pulse.

For the PIF neuron case, the corresponding SDC formula can be directly obtained from Eqs. (24) and (25) by turning there *τ*_*m*_ → +∞ and *I*_*c*_ → 0, with *I*_*c*_*τ*_*m*_ = *Q*_*c*_ (see details in Sec. 2). In particular, denoting *τ*_*a*_*/τ*_*m*_ = *z* → 0, from Eq. (25) we asymptotically get *K* = −*W*_−1_(−*z* exp(−*z*)), which is equivalent to −*K* exp(−*K*) = −*z* exp(−*z*), according to the Lambert W function definition. Substituting the latter expression into Eq. (24), we get

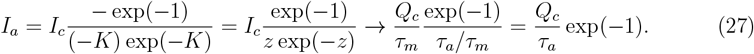

This result is exact for the PIF neuron and coincides with the classic SDC fitting by a hyperbola.

## 5. Comparison with other pulse shapes

It seems advisable to compare the obtained SDC formula with the results for (i) a standard rectangular pulse, (ii) linearly-increasing ramp pulse, and (iii) instantly rising and exponentially decaying pulse. For the first two shapes, parameter *τ*_*a*_ stands for the pulse duration, and, for the latter, *τ*_*a*_ is the time constant of exponential decay.

1. For the rectangular pulse

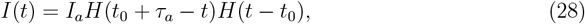

where *H*(*x*) is the Heaviside unit-step function

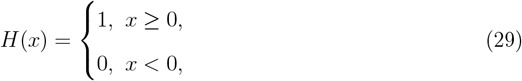

the first spike time *t*_*sp*_ for the LIF neuron (at *t*_0_ = 0 and *V* (*t*_0_) = *V*_*rest*_) is given by Eq. (3), where *τ*_*ref*_ = 0,

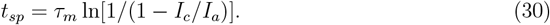

The spike must occur before the end of the stimulating pulse, i.e. *t*_*sp*_ ≤ *τ*_*a*_. The limiting equality *t*_*sp*_ = *τ*_*a*_ determines the SDC,

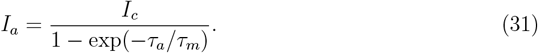

It is widely accepted that this formula was first reported by L. Lapicque in 1907 [15] (cp. [16]). Limiting cases *τ*_*a*_ ≫ *τ*_*m*_ and *τ*_*a*_ ≪ *τ*_*m*_ for Eq. (31) give *I*_*a*_ ≈ *I*_*c*_ and *I*_*a*_ ≈ *I*_*c*_*τ*_*m*_*/τ*_*a*_, respectively. For the PIF neuron, Eq. (31) is transformed to *I*_*a*_ = *Q*_*c*_*/τ*_*a*_.

2. For the ramp pulse given by

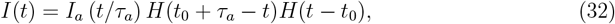

with the same initial conditions (*t*_0_ = 0, *V* (*t*_0_) = *V*_*rest*_) one gets the first spike condition for the LIF neuron

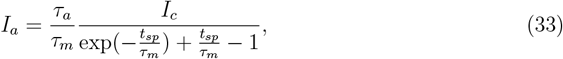

where, as previously, *t*_*sp*_ ≤ *τ*_*a*_, and the SDC is determined by *t*_*sp*_ = *τ*_*a*_, resulting in

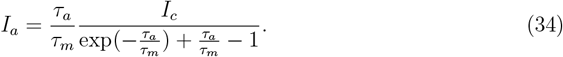

Limiting cases *τ*_*a*_ ≫ *τ*_*m*_ and *τ*_*a*_ ≪ *τ*_*m*_ for Eq. (34) give *I*_*a*_ ≈ *I*_*c*_ and *I*_*a*_ ≈ 2*I*_*c*_*τ*_*m*_*/τ*_*a*_, respectively. For the PIF neuron, Eq. (34) is transformed to *I*_*a*_ = 2*Q*_*c*_*/τ*_*a*_.

3. Finally, for the instantly rising and exponentially decaying pulse,

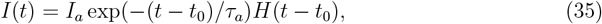

from Eqs. (8) we get

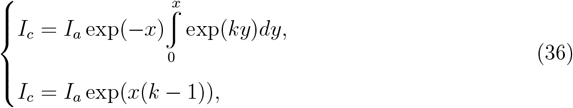

where, as before, *x* = *t*_*sp*_*/τ*_*m*_ and *k* = 1 − *τ*_*m*_*/τ*_*a*_. If *k* = 0, the solution of Eqs. (36) is *x* = 1 and *I*_*a*_ = *I*_*c*_ exp(1). If *k* ≠ 0,

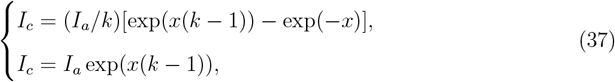

from where one gets *k* = 1 − (*I*_*c*_*/I*_*a*_)^*k/*(1−*k*)^ and *I*_*a*_ = *I*_*c*_(1 − *k*)^−(1−*k*)*/k*^. In usual notations, the SDC reads (cp. [19])

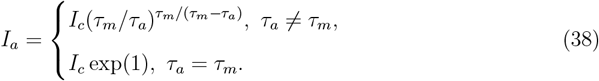

At *τ*_*a*_ → +∞, taking into account that

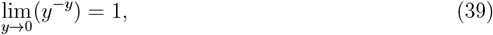

where *y* = *τ*_*m*_*/τ*_*a*_, one gets the standard value *I*_*a*_(*τ*_*a*_ → +∞) = *I*_*c*_. The case *τ*_*a*_ ≪ *τ*_*m*_ for Eq. (38) gives 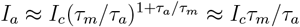. For the PIF neuron, Eq. (38) is transformed to *I*_*a*_ = *Q*_*c*_*/τ*_*a*_, which is exactly the same as for the rectangular pulse case.

Summarizing, we have explicitly shown that different pulse shapes lead to significant functional differences in the SDC for the LIF neuron, however, up to a numerical factor, they are completely insignificant for the case of the PIF neuron, which has a relatively universal SDC in the form of a hyperbola. That universality can be explained as follows: for the PIF neuron the SDC comes from the condition that, for initiating a spike, the total electric charge *Q*_*tot*_ delivered by the incoming excitatory pulse, i.e.

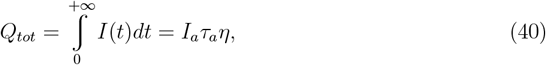

where *η* is a numerical constant determined by the pulse shape, should be equal to the critical charge *Q*_*c*_ (see Sec. 2). All the above results for the SDC in the PIF neuron case follow directly from equality *Q*_*tot*_ = *Q*_*c*_.

## 6. Conclusion

For the LIF neuron stimulated by an excitatory current pulse in the form of the alpha function, analytical strength-duration curve (SDC) for the spiking response has been accurately found. The SDC is expressed through the Lambert W function, a well-studied and widely-implemented special function. In addition, general SDC equations for an arbitrary shape of the stimulating pulse have been obtained. For comparison, we have also re-derived in a unified way the analytical SDCs for (i) a standard rectangular pulse, (ii) ascending ramp pulse, and (iii) instantly rising and exponentially decaying pulse. In the limit of no leakage, it has been shown that for all considered cases the SDC is reduced to the classical hyperbola.

## Data and code availability

The Supplementary Material for this article contains (i) the data for graphs in Figure 1 and (ii) ready-to-use MATLAB codes for reproducing the data. It is available online at https://doi.org/10.6084/m9.figshare.24081849.

## Acknowledgments

The author thanks T.S. Zemskova and N.D. Efimova for verifying some of the results. This work was supported by a European Research Council Consolidator Grant (SYNAPSEEK, 819603, to Tim P. Vogels).

## References

[1] L.F. Abbott, C. van Vreeswijk, Asynchronous states in networks of pulse-coupled oscillators, Phys. Rev. E 48, 1483–1490 (1993). 10.1103/PhysRevE.48.1483

[2] D. Hansel, G. Mato, C. Meunier, Synchrony in excitatory neural networks, Neural Comput. 7, 307–337 (1995). 10.1162/neco.1995.7.2.307

[3] T.P. Vogels, L.F. Abbott, Signal propagation and logic gating in networks of integrate-and-fire neurons, J. Neurosci. 25, 10786–10795 (2005). 10.1523/jneurosci.3508-05.2005

[4] M. Sahin, Y. Tie, Non-rectangular waveforms for neural stimulation with practical electrodes, J. Neural Eng. 4, 227–233 (2007). 10.1088/1741-2560/4/3/008

[5] T.J. Foutz, C.C. McIntyre, Evaluation of novel stimulus waveforms for deep brain stimulation, J. Neural Eng. 7, 066008 (2010). 10.1088/1741-2560/7/6/066008

[6] A. Wongsarnpigoon, J.P. Woock, W.M. Grill, Efficiency analysis of waveform shape for electrical excitation of nerve fibers, IEEE Trans. Neural Syst. Rehabil. Eng. 18, 319–328 (2010). 10.1109/TNSRE.2010.2047610

[7] J.-I. Lee, M. Im, Non-rectangular waveforms are more charge-efficient than rectangular one in eliciting network-mediated responses of ON type retinal ganglion cells, J. Neural Eng. 15, 055004 (2018). 10.1088/1741-2552/aad416

[8] A.V. Hill, Excitation and accommodation in nerve, Proc. R. Soc. B 119, 305–355 (1936). 10.1098/rspb.1936.0012

[9] D. Noble, R.B. Stein, The threshold conditions for initiation of action potentials by excitable cells, J. Physiol. 187, 129–162 (1966). 10.1113/jphysiol.1966.sp008079

[10] B.I. Khodorov, The Problem of Excitability (Plenum Press, 1974). 10.1007/978-1-4613-4487-2

[11] J.J.B. Jack, D. Noble, R.W. Tsien, Electric current flow in excitable cells (Clarendon Press, Oxford, 1975).

[12] R.J. MacGregor, E.R. Lewis, Neural Modeling: Electrical Signal Processing in the Nervous System (Plenum Press, 1977). 10.1007/978-1-4684-2190-3

[13] H.C. Tuckwell, Introduction to Theoretical Neurobiology. Volume 1: Linear Cable Theory and Dendritic Structure (Cambridge University Press, 1988). 10.1017/CBO9780511623271

[14] E.J. Tehovnik et al., Direct and indirect activation of cortical neurons by electrical microstimulation, J. Neurophysiol. 96, 512–521 (2006). 10.1152/jn.00126.2006

[15] N. Brunel, M.C.W. van Rossum, Quantitative investigations of electrical nerve excitation treated as polarization, Biol. Cybern. 97, 341–349 (2007). 10.1007/s00422-007-0189-6

[16] J.L. Hoorweg, Ueber die elektrische Nervenerregung, Pflügers Arch. 52, 87–108 (1892). 10.1007/BF01661875

[17] G. Weiss, Sur la possibilite de rendre comparables entre eux les appareils servant a l’excitation electrique, Arch. Ital. Biol. 35, 413–446 (1901). https://architalbiol.org/index.php/aib/article/view/35413

[18] A.V. Hill, A new mathematical treatment of changes of ionic concentration in muscle and nerve under the action of electric currents, with a theory as to their mode of excitation, J. Physiol. 40, 190–224 (1910). 10.1113/jphysiol.1910.sp001366

[19] H.A. Blair, On the intensity-time relations for stimulation by electric currents. II. J. Gen. Physiol. 15, 731–755 (1932). 10.1085/jgp.15.6.731

[20] T.H. Brown, D. Johnston, Voltage-clamp analysis of mossy fiber synaptic input to hippocampal neurons, J. Neurophysiol. 50, 487–507 (1983). 10.1152/jn.1983.50.2.487

[21] S.H. Williams, D. Johnston, Kinetic properties of two anatomically distinct excitatory synapses in hippocampal CA3 pyramidal neurons, J. Neurophysiol. 66, 1010–1020 (1991). 10.1152/jn.1991.66.3.1010

[22] R.M. Corless et al., On the Lambert W function, Adv. Comput. Math. 5, 329–359 (1996). 10.1007/BF02124750

[23] J.P. Boyd, Solving Transcendental Equations (Society for Industrial and Applied Mathematics, 2014). 10.1137/1.9781611973525

[24] A.N. Burkitt, G.M. Clark, Analysis of integrate-and-fire neurons: synchronization of synaptic input and spike output, Neural Comput. 11, 871–901 (1999). 10.1162/089976699300016485

[25] W.H. Nesse, A. Borisyuk, P.C. Bressloff, Fluctuation-driven rhythmogenesis in an excitatory neuronal network with slow adaptation, J. Comput. Neurosci. 25, 317–333 (2008). 10.1007/s10827-008-0081-y

[26] E. Nordlie, T. Tetzlaff, G.T. Einevoll, Rate dynamics of leaky integrate-and-fire neurons with strong synapses, Front. Comput. Neurosci. 4, 149 (2010). 10.3389/fncom.2010.00149

[27] B. Kriener et al., Dynamics of self-sustained asynchronous-irregular activity in random networks of spiking neurons with strong synapses, Front. Comput. Neurosci. 8, 136 (2014). 10.3389/fncom.2014.00136

[28] S. Ashhad et al., Microcircuit synchronization and heavy-tailed synaptic weight distribution augment preBötzinger Complex bursting dynamics, J. Neurosci. 43, 240–260 (2023). 10.1523/jneurosci.1195-22.2022

[29] A. Morrison, M. Diesmann, W. Gerstner, Phenomenological models of synaptic plasticity based on spike timing, Biol. Cybern. 98, 459–478 (2008). 10.1007/s00422-008-0233-1

[30] A. Destexhe, Z.F. Mainen, T.J. Sejnowski, Synthesis of models for excitable membranes, synaptic transmission and neuromodulation using a common kinetic formalism, J. Comput. Neurosci. 1, 195–230 (1994). 10.1007/BF00961734

[31] J. Göltz et al., Fast and energy-efficient neuromorphic deep learning with first-spike times, Nat. Mach. Intell. 3, 823–835 (2021). 10.1038/s42256-021-00388-x

[32] I.-M. Comsa et al., Temporal coding in spiking neural networks with alpha synaptic function: learning with backpropagation, IEEE Trans. Neural Netw. Learn. Syst. 33, 5939–5952 (2022). 10.1109/tnnls.2021.3071976

